# Is Inappropriate Pulse Timing Responsible for Poor Binaural Hearing with Cochlear Implants?

**DOI:** 10.1101/2023.08.04.551983

**Authors:** Jan W. H. Schnupp, Sarah Buchholz, Alexa N. Buck, Henrike Budig, Lakshay Khurana, Nicole Rosskothen-Kuhl

## Abstract

Cochlear implants (CIs) have restored enough of a sense of hearing to around one million severely hearing impaired patients to enable speech understanding in quiet. However, several aspects of hearing with CIs remain very poor. This includes a severely limited ability of CI patients to make use of interaural time difference (ITD) cues for spatial hearing and noise reduction. A major cause for this poor ITD sensitivity could be that current clinical devices fail to deliver ITD information in a manner that is accessible to the auditory pathway. CI processors measure the envelopes of incoming sounds and then stimulate the auditory nerve with electrical pulse trains which are amplitude modulated to reflect incoming sound envelopes. The timing of the pulses generated by the devices is largely or entirely independent of the incoming sounds. Consequently, bilateral CIs (biCIs) provide veridical envelope (ENV) ITDs but largely or entirely replace the “fine structure” ITDs that naturally occur in sounds with completely arbitrary electrical pulse timing (PT) ITDs. To assess the extent to which this matters, we devised experiments that measured the sensitivity of deafened rats to precisely and independently controlled PT and ENV ITDs for a variety of different CI pulse rates and envelope shapes. We observed that PT ITDs completely dominate ITD perception, while the sensitivity to ENV ITDs was almost negligible in comparison. This strongly suggests that the confusing yet powerful PT ITDs that contemporary clinical devices deliver to biCI patients may be a major cause of poor binaural hearing outcomes with biCIs.

**Significance Statement:** CIs deliver spectro-temporal envelopes, including speech formants, to severely deaf patients, but they do little to cater to the brain’s ability to process temporal sound features with sub-millisecond precision. CIs “sample” sound envelope signals rapidly and accurately, and thus provide information which should make it possible in principle for CI listeners to detect envelope ITDs in a similar way to normal listeners. However, here we demonstrate through behavioral experiments on CI implanted rats trained to detect sub-millisecond ITDs that pulse timing ITDs completely dominate binaural hearing. This provides the strongest confirmation to date that the arbitrary pulse timing widely used in current clinical CIs is a critical obstacle to good binaural hearing through prosthetic devices.

## Introduction

The human brain is normally remarkably good at locating sound sources in space, which also assists in our ability to separate foreground from background sounds. This type of directional hearing remains, however, largely out of reach for severely hearing impaired patients who rely on cochlear implants (CIs) for their perception of the auditory world. This is due, at least in large part, to their dramatically reduced sensitivity to an important sound source location cue: interaural time differences (ITDs) (Kerber and Seeber 2012; Litovsky and Gordon 2016; Ehlers et al. 2017).

Originally, severely hearing impaired patients were fitted with only a single CI, as restoring monaural hearing is often sufficient to enable speech understanding, at least in quiet. However, good spatial hearing, as well as target source separation, relies heavily on binaural hearing. Nowadays, patients are increasingly fitted with CIs in both ears, which can bring significant improvements in speech in noise detection thresholds, and even some sound localization due to an ability to detect interaural differences in sound intensity (also known as interaural level differences or ILDs) (Gordon et al. 2014; Sparreboom et al. 2022). Nevertheless, bilaterally supplied CI patients typically remain largely insensitive to ITDs, unlike normally hearing (NH) humans or many other mammals who are exquisitely sensitive to the minuscule ITDs that arise when a sound wave arriving from one side strikes the near ear a fraction of a millisecond before reaching the far ear. Natural hearing affords typical ITD detection thresholds as low as 10 to 20 μs (Klumpp and Eady 1956; Zwislocki and Feldman 1956; Brughera et al. 2013). In contrast, CI patients typically have no ITD sensitivity at all when tested with their standard issue clinical devices running at high stimulation rates. They usually need to be tested with specialized experimental processors at pulse rates ≤100 pps to demonstrate any measurable ITD sensitivity, and even then their thresholds are rarely better than a few hundred μs, and, at the higher pulse rates required to allow effective speech encoding, they are often much worse (Litovsky et al. 2012; Laback et al. 2015). Prelingually deaf CI patients are particularly affected, and only rarely show any measurable ITD sensitivity even when tested with experimental processors at low stimulation rates (Litovsky et al. 2010; Ehlers et al. 2017).

While the reasons behind the poor ITD sensitivity of bilateral CI (biCI) patients are still actively debated, recent studies on rats have demonstrated that such poor ITD sensitivity may not be inevitable. Indeed, neonatally deafened (ND) rats implanted with CIs in early adulthood can achieve excellent ITD sensitivity if they are provided with temporally precise electric pulses straight after implantation (Rosskothen-Kuhl et al. 2021; Buck et al. 2023). This discovery not only indicates that the very poor ITD sensitivity routinely exhibited by CI patients is very likely preventable, it also makes it possible for the first time to investigate experimentally which stimulation parameters and strategies are needed to permit excellent ITD sensitivity in prosthetic hearing. One important question which we address here is whether ITD cues have to be delivered in the timing of electrical pulses to make good ITD sensitivity possible, or whether ITDs could plausibly alternatively be delivered in the envelope of pulse trains. If delivering ITDs in pulse timing turns out to be essential, then this has important implications. Not only would it provide a likely explanation for why ITD perception is so poor with current biCIs, but it would also provide a strong argument that the CI processors currently issued to biCI patients may need a substantial redesign.

Essentially all of the about 1 million CI devices in current clinical use processing strategies, which stimulate the auditory nerve (AN) with trains of electrical pulses that are amplitude modulated such that the pulse amplitudes reflect the envelopes of incoming sounds. The timing of the individual pulses in the pulse trains is usually set by the internal clock of the CI processor, and, depending on manufacturer and model, is either largely or entirely independent of temporal features of the sounds that are encoded. Thus little to none of the temporal fine structure (FS) of incoming sounds is encoded in the pulse timing (PT) of the electrical stimuli, and the devices rely largely or entirely on encoding sound envelopes (ENV) in the pulse amplitude modulation (Carlyon and Goehring 2021) to encode sounds. As a consequence, biCI users will normally receive ENV ITDs that veridically encode binaural cues, but arbitrary PT ITDs which provide no useful binaural cue information. In fact, PT ITDs may instead be a significant source of confusion. The extent to which this potential confusion matters depends on the relative sensitivity of the auditory pathway to PT and ENV ITDs, respectively. If the CI stimulated brain is sensitive to PT but not ENV ITDs, then that might explain the poor ITD discrimination performance typically observed in biCI users.

What complicates this picture is that, when biCI patients tested with experimental processors that permit the delivery of precisely controlled PT ITDs still tend to perform very poorly. Indeed, most of the studies mentioned above were carried out with such experimental processors. But it is very hard to know what one should conclude from the fact that poor ITD sensitivity persists in biCI patients when the arbitrary PT ITDs normally delivered by clinical devices are replaced by accurate PT ITDs for the relatively short period of a psychoacoustic experiment. It is well known that CI users require adaptation periods extending over many months for their auditory pathways to adapt to the rather unnatural input provided by their devices before they can make effective use of their input (Tyler et al. 1997). It is reasonable to expect that the neural pathways of these patients substantially reorganizes during this adaptation period, presumably in a manner that makes informative features of the provided input more salient by reducing the sensitivity to uninformative features of the input, such as the arbitrary timing of the pulses emitted by their clinical processors. When such patients are then tested acutely for PT and ENV ITD sensitivity after months of stimulation with input in which PT was at best useless and at worst confusing, the observed thresholds (Noel and Eddington 2013) are unlikely to be representative of the system’s underlying capability. To understand the true potential of PT ITDs and ENV ITDs, respectively, to either inform or confuse a biCI recipient’s brain, one needs to study their influence in participants whose auditory pathway has not undergone months of training and experience listening to the world through devices which effectively disincentivize PT ITD sensitivity. However, despite their limitations, current clinical processors do deliver undeniable benefits, and in the absence of clearly better alternatives, clinical best practice dictates that patients should receive such devices and use them as much as possible in their daily lives. In the present context, clinical, ethical, and practical considerations therefore prevent experiments on human patients which would shed a clear light on the true potential of PT and ENV ITDs, respectively, in the CI stimulated mammalian pathway. We therefore turned to work in experimental animals to address the important question whether ITD cues have to be delivered in the timing of electrical pulses or can alternatively be delivered in the envelope of pulse trains to enable good ITD sensitivity. For this purpose, we conducted a series of behavioral experiments performed on eight ND rats, which were implanted with biCIs in young adulthood and trained and tested from the beginning in a stimulus lateralization task (Fig. 1A) with precisely controlled combinations of PT ITD and ENV ITD. This enabled us to elucidate the relative effectiveness of these two types of binaural cues in informing bionic spatial hearing.

**Figure 1:**
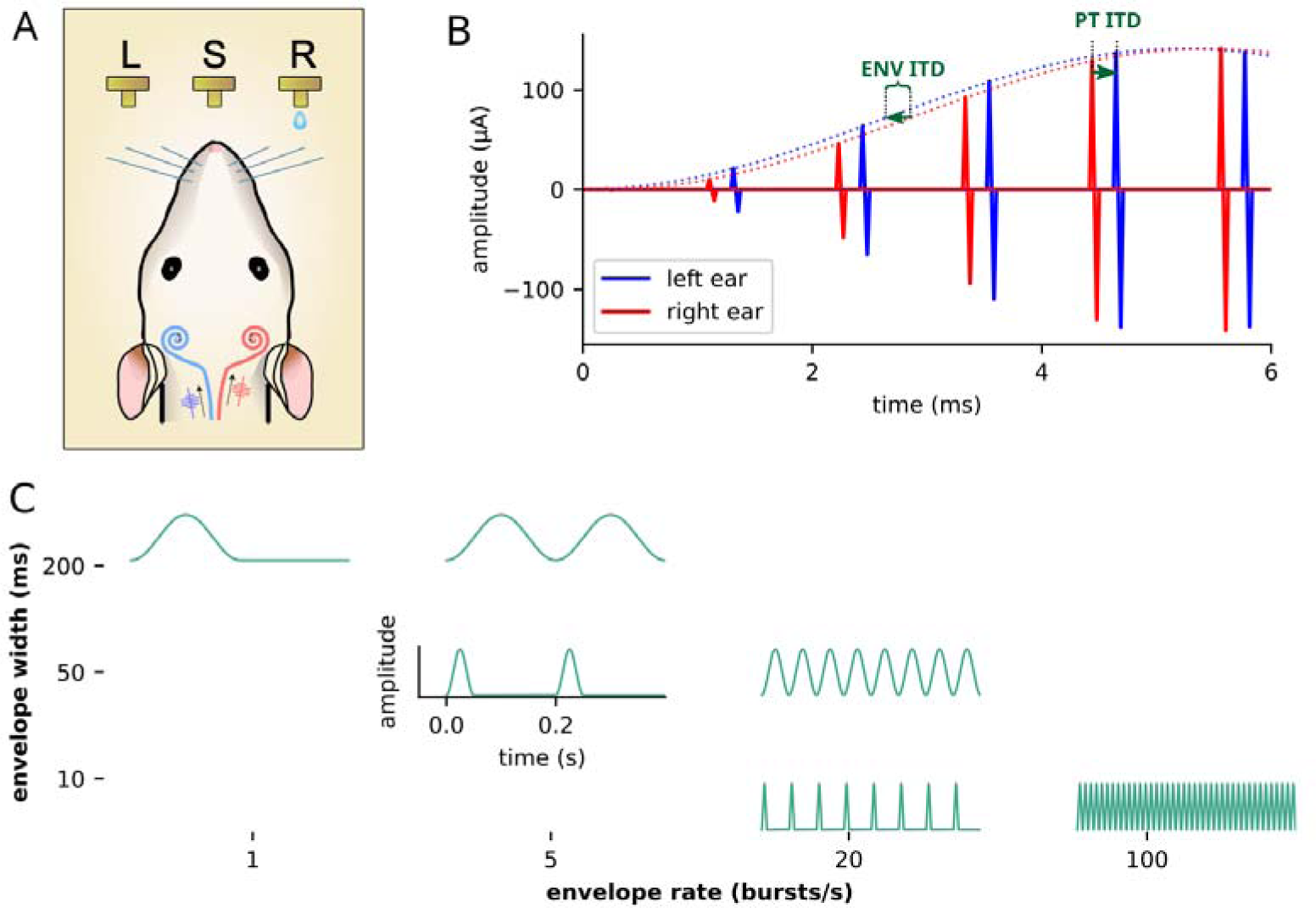
A) Cartoon illustrating the experimental procedure. ND rats were fitted with biCIs and trained to initiate trials by licking a central start spout (“S”). This triggered electric pulse trains to be delivered, with a small ITD, to their left (blue) and right (red) ears. To receive a water reward, the animals had to lick response spouts on the side (left “L”, or right “R”) corresponding to the side of the earlier pulse train. In this example the ride ear is leading. B) Schematic of the types of binaural pulse trains used in this experiment. Hanning window envelopes were used to amplitude modulate biphasic pulse trains. The ITD of the envelopes (ENV ITD) could be varied independently of the pulse timing ITD (PT ITD). In the example shown, the left ear (blue) envelope rises earlier than the right ear (red) envelope, but the left ear pulses occur after the right ear pulses. C) Schematic illustrating the six envelope types tested. To address the possibility that ENV ITD sensitivity might depend on the shape of the envelope, six different envelope types were tested, differing in envelope width (200, 50, or 10 ms) and presentation with or without silent gaps (envelope rate either equal to or greater than 1/envelope width).

## Results

Eight ND rats received chronic biCIs in early adulthood (postnatal days 70-119) and were successfully trained to lateralize ITD cues within 3-5 days of behavioral training. Figure 1 shows the experimental procedure and the different stimulation parameters tested. In short, the rats learned to lick a central “start” spout (S) to trigger a brief binaural pulse train to their CIs with an ITD of either −80 μs or +80 μs (here we use negative numbers to denote left ear leading ITDs). We focused deliberately on ITDs of +/-80 μs because ITDs of that magnitude are very easy to discriminate for NH humans as well as for NH and biCI rats (Li et al. 2019; Rosskothen-Kuhl et al. 2021), but they are impossibly hard to lateralize for human biCI patients. Improving outcomes for severely hearing impaired patients will require that we understand what makes lateralizing such moderately small ITDs difficult in the biCI context.

The rats were required to choose one of two response spouts, either on the left (L) or on the right (R; Fig. 1A). If they licked on the side of the leading ear, their choice was reinforced with a drinking water reward. Incorrect choices elicited negative feedback in the form of a short timeout. During initial training, the animals were given exclusively stimuli in which PT ITD and ENV ITD always agreed, so there was no ambiguity about which side would be a correct response deserving of a reward. After the initial training, we introduced stimuli in which PT and ENV ITDs were allowed to vary independently, including “probe trials” (1 in 6 trials chosen at random), in which PT ITD and ENV ITD values pointed in opposite directions (Fig. 1B). In such probe trials, the animals were rewarded irrespective of the spouts they chose. By analyzing the animals’ responses to a very large number of stimuli with different PT and ENV ITD combinations, we were then able to quantify how strongly PT ITDs and ENV ITDs, respectively, can inform sound location judgments in the biCI stimulated auditory system. For more details on the behavioral training and test setup see Rosskothen-Kuhl et al. (2021) and Methods below.

We needed to bear in mind that the effectiveness of ENV ITDs may depend on parameters such as envelope shape and pulse rate. For example, it may be easier for the auditory brain to process ENV ITDs when stimulus envelopes have rapidly rising slopes which are densely sampled by pulses occurring at a high rate, than if envelope slopes are shallow and thus spread out in time. Indeed, recordings in the inferior colliculus by Smith and Delgutte (2008) support this idea. Similarly, ENV ITDs might be less effective if their shape is only sparsely sampled by pulses occurring at a low rate. We therefore tested a wide range of stimulus envelopes, from quite slow to very rapid (Fig. 1C), and we also tested envelopes with or without silent gaps, as fast attacks and gaps have been shown to be of potential importance in ENV ITD sensitivity in normal hearing, both psychophysically (Klein-Hennig et al. 2011) and in IC recordings in anesthetized guinea pigs (Greenberg et al. 2017). These were presented at two pulse rates, 900 pps, a very commonly used rate in clinical settings, and 4500 pps, which is very fast and might favor ENV ITD encoding.

To visualize how stimulus ITD influenced the animals’ lateralization choice, we pooled the data from all rats and calculated the proportion of “right” responses (that is, the proportion of trials in which the animals chose the right response spout) for each of the tested PT ITD and ENV ITD combinations. The result of this analysis, pooled over all envelope types but separated out for the two different pulse rates, is shown in 3D bar charts in Figure 2. It can be seen that at a clinical pulse rate of 900 pps (Fig. 2, left), a change in PT ITDs strongly influenced the lateralization decision of our CI rats, recognizable by an average of ∼87% responses to “right” and orange bars when the PT ITD was +80 µs (right ear leading) and only ∼16.8% and purple bars when the PT ITD was −80 µs (left ear leading). In contrast, changing the ENV ITD had a much smaller effect on the rats’ decision. Particularly for non-zero PT ITDs, % “right” responses barely change at all with changing ENV ITD. Contrary to our hypothesis above, increasing the pulse rate from 900 to 4500 pps had no obvious positive effect on ENV ITD sensitivity of our CI rats. In fact, the lateralization decision of the animals is still dominated by the PT ITD, even if the effect of PT ITD was somewhat smaller than at 900 pps (Fig. 2, right).

**Figure 2:**
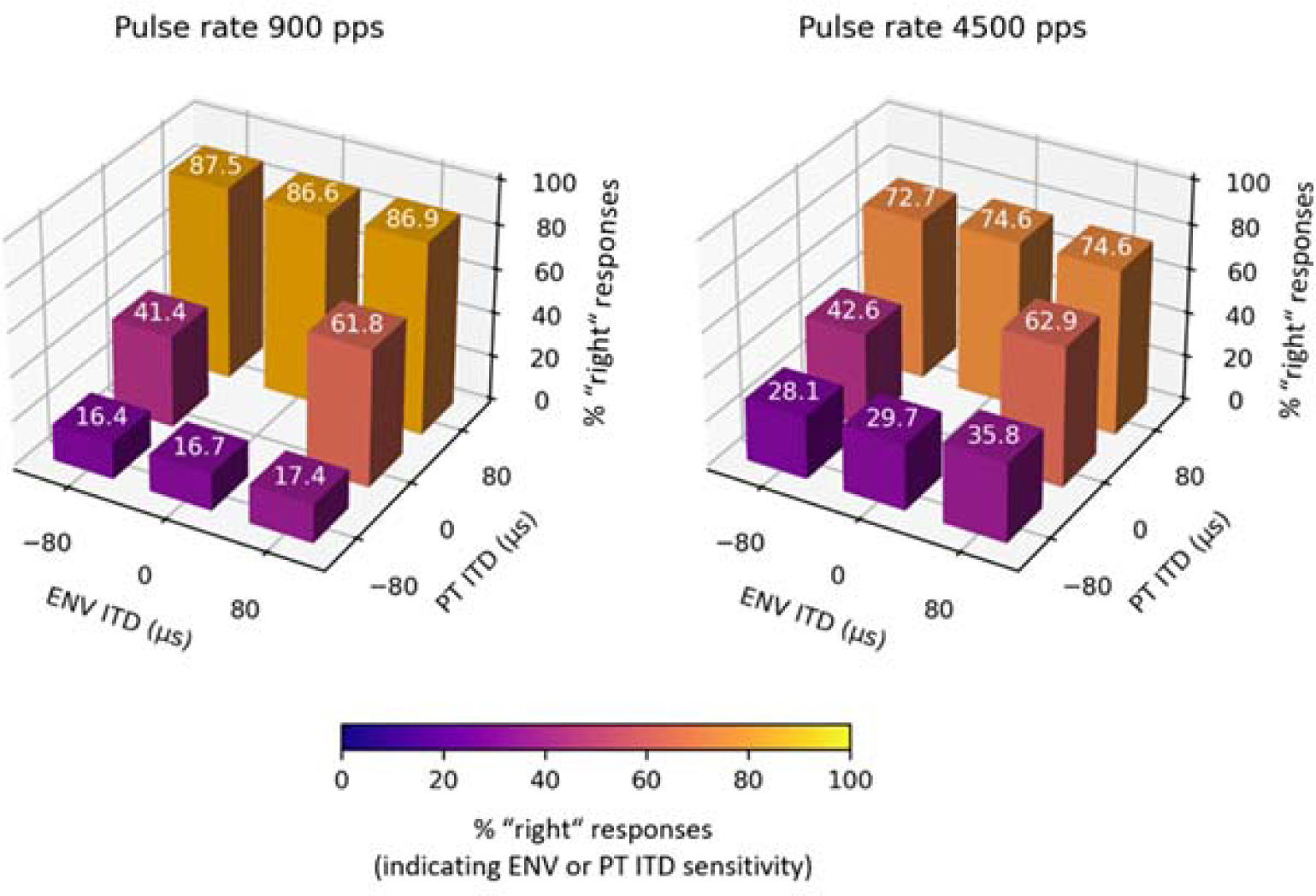
3D bar charts showing the summarized proportion of “right” responses for each of the two different stimulus pulse rates (left: 900 pps; right: 4500 pps). The numbers in white above each bar give the percentage of “right” responses for stimuli with PT ITD and ENV ITD values given by each panel’s x-axis and y-axis coordinate. The color of each bar also indicates the percent of responses on the right hand side according to the colorbar shown at the bottom. Percent “right” responses increased very substantially as PT ITDs changed from −80 µs (left leading ear) to +80 μs (right leading ear). Changes in “right” responses as a function of ENV ITD are much less pronounced.

Figure 3 shows the effect of the different envelope types and gaps (Fig. 1C) on the ITD sensitivity of our CI rats. In each panel responses to different PT ITDs are arranged bottom to top, while ENV ITDs are arranged left to right. The color scale (see color bar bottom right) makes the effect of PT ITD and ENV ITD, respectively, visible as a color gradient. PT ITD clearly has a powerful influence, as all panels exhibit a very strong vertical color gradient. This is most pronounced for stimuli at 900 pps (Fig. 3, top row). In the best case (top-right), the responses increased from only 10% or less “right” responses for PT ITDs of −80 μs (left leading ear) to 90% or more for PT ITDs of +80 μs (right leading ear). In Figure 3, effects of ENV ITD should manifest as horizontal color gradients. These are much less pronounced than the strong and consistent vertical PT ITD-associated gradients, but they are not entirely absent. For every pulse rate and envelope type tested, there were fewer “right” responses for (−80 μs ENV ITD, 0 μs PT ITD) stimuli than for (+80 μs ENV ITD, 0 μs PT ITD) stimuli. Given the very large number of trials in this dataset (altogether 16540 trials with PT ITD=0) this is very unlikely to be a coincidence, and statistical analysis described next confirmed that the effect of ENV ITD, albeit small, was nevertheless greater than zero in many of the stimulus conditions tested.

**Figure 3:**
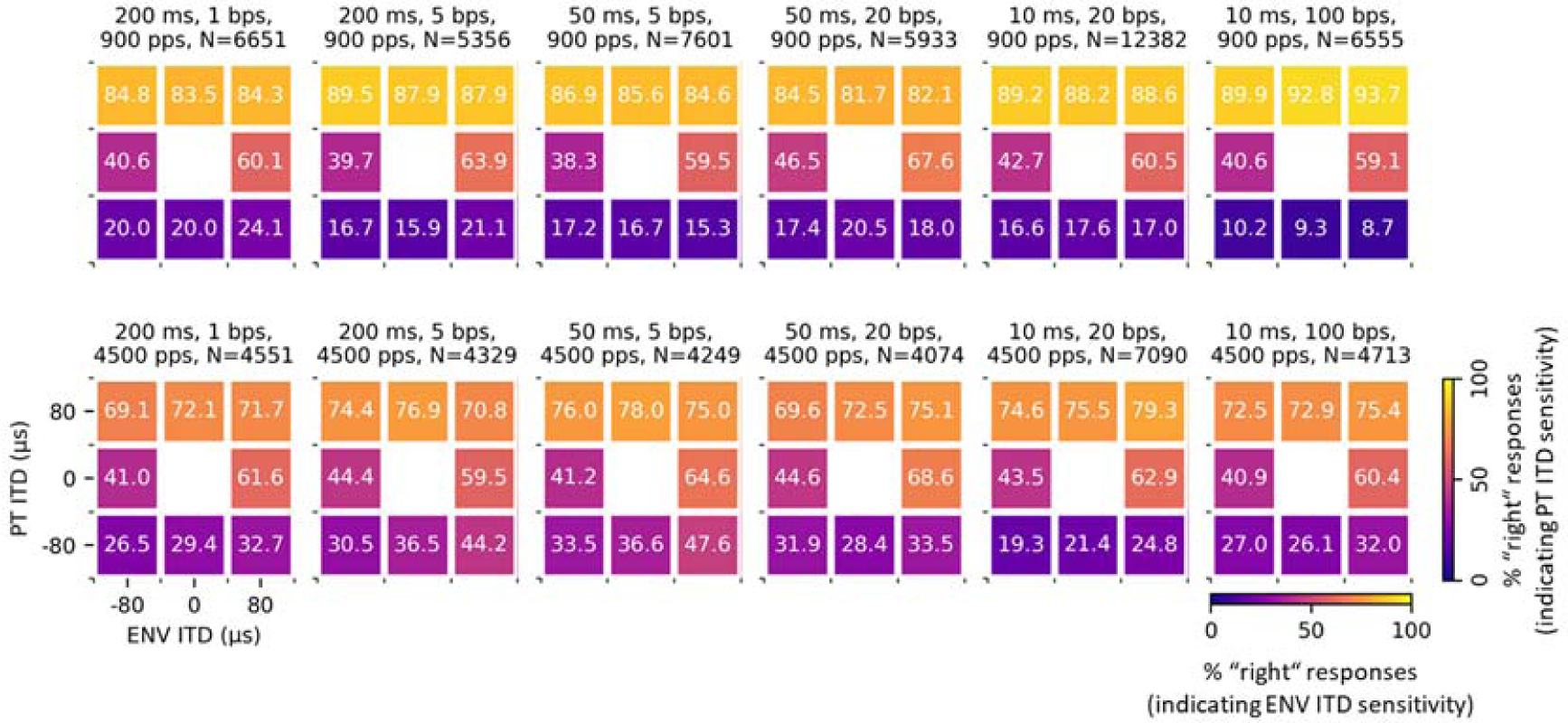
Heatmaps showing the proportion of “right” responses for each of the two different stimulus pulse rates (rows) and the six different envelope types shown in Fig. 1C (columns). The headings above each panel indicate the envelope type, the pulse rate (pps), and the total number of trials (N), pooled across all eight animals, for each condition. The numbers in white in each cell give the percentage of “right” responses for stimuli with PT ITD and ENV ITD values given by each cell’s row and column coordinate, as indicated at the bottom left. The color of each cell also indicates the percent “right” responses according to the colorbars shown at the bottom right. Percent “right” responses increased very substantially as PT ITDs changed from −80 µs (left leading ear) to +80 μs (right leading ear). Changes in “right” responses as a function of ENV ITD are much less pronounced.

To quantify the effects of PT ITD and ENV ITD observed in Figure 3, we fitted the behavioral data with a probit regression model described in detail in the Methods section below. The probit coefficients for PT ITD and ENV ITD returned by the probit models are somewhat akin to regression slopes. Values near zero indicate small or negligible effects of the corresponding ITD type, while large positive values indicate a strong, monotonic increase in the proportion of “right” responses with increasing ITD. Separate probit models were fitted for each pulse rate, envelope type, and animal. The coefficients obtained are summarized as bar charts in Figure 4.

**Figure 4:**
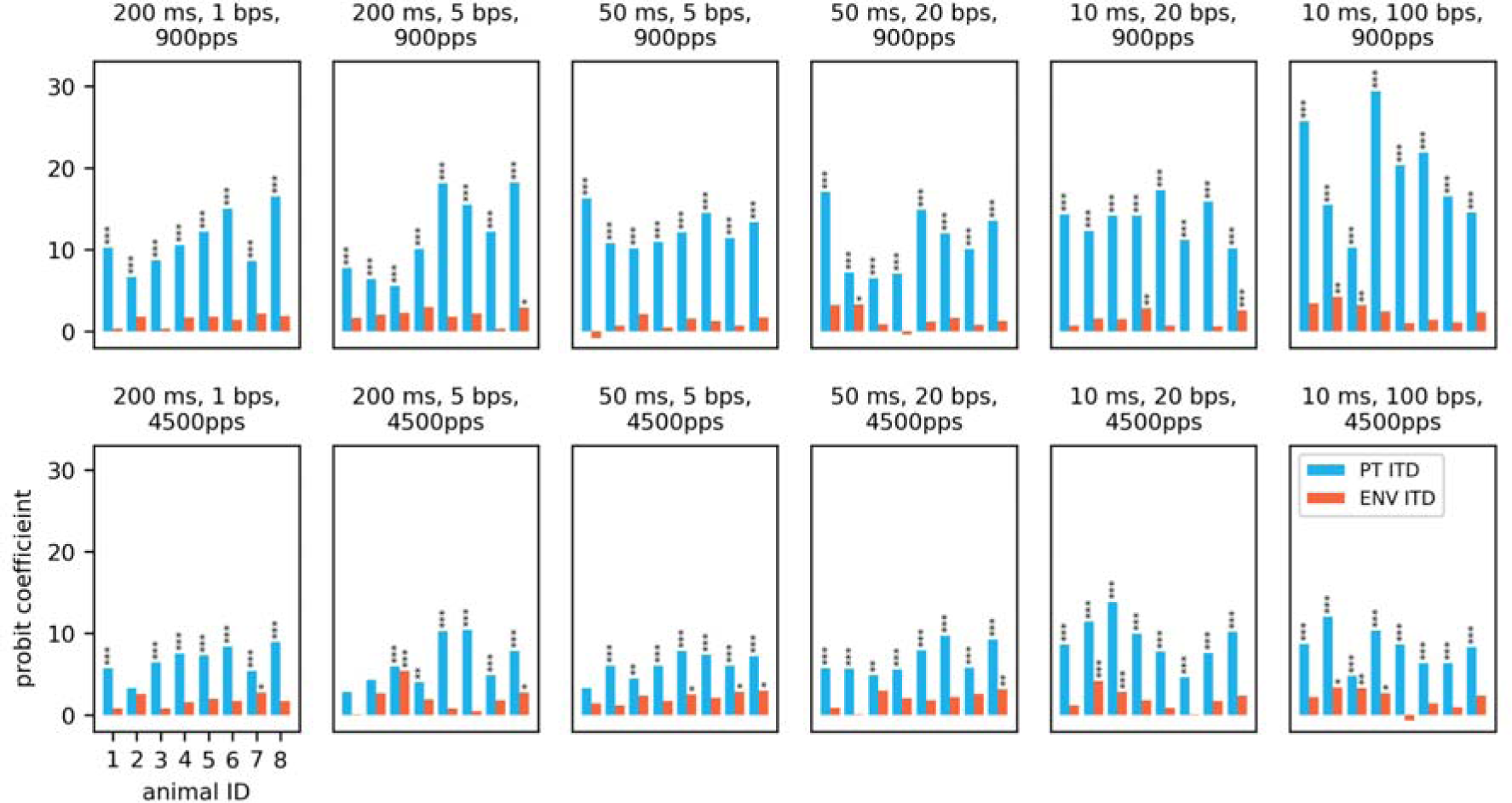
Behavioral sensitivity to PT ITD (blue bars) and ENV ITD (red bars) quantified as probit coefficients for each of the eight animals. Each panel shows probit coefficients for a different stimulus envelope type and pulse rate, as indicated above each panel. The small asterisks indicate which probit coefficients were statistically greater than zero at p<0.05 (*), p<0.01 (**) or p<0.001 (***), respectively (Bonferroni corrected).

As shown in Figure 4, the PT ITD coefficients were significant in almost all cases, and seemed to vary as a function of stimulus condition, from a minimum (averaged over the eight rats) of 6.03 for 4500 pps stimuli with 50 ms wide envelopes at 5 bursts/s (bps), to a maximum of 19.23 for 900 pps stimuli with 10 ms wide envelopes at 100 bps. In contrast, ENV ITD coefficients were much smaller, they failed to reach statistical significance in many individual cases, and they varied much less as a function of stimulus condition, from a minimum of 0.95 for 900 pps stimuli with 50 ms wide envelopes at 5 bps, to a maximum of 2.403 for 900 pps stimuli with 10 ms wide envelopes at 100 bps. Note, however, that although ENV ITDs coefficients often failed to reach significance for any one animal in any one condition, only 4 of the 96 ENV ITDs coefficients shown in Figure 4 were below zero. If ENV ITD had no effect overall, we would expect negative coefficients to be as common as positive ones. We can therefore conclude that ENV ITD did have an effect, even if that effect is so small that data from a single animal at a single condition was usually not sufficient to demonstrate it.

In comparison, we note that PT ITD coefficients were on average 6.2 times larger than ENV ITD coefficients, with ratios ranging from a minimum of 2.8 times larger for (4500 pps, 50 ms, 5 bps) stimuli to a maximum of 13.07 times larger for (900 pps, 50 ms, 5 bps) stimuli. Finally, we note that the results were quite consistent across all eight animals in our study, with all animals achieving PT ITD coefficients of 10 or above for at least some stimulus conditions, and all animals showing similar trends, with on average larger PT ITD coefficients at 900 pps compared to 4500 pps, and considerably smaller ENV ITD than PT ITD coefficients throughout.

To further investigate the effect of pulse rate and envelope type on the probit coefficients shown in Figure 4, we computed a two-way repeated measures ANOVA using the pingouin library (https://pingouin-stats.org). For PT ITD (Fig. 4, blue bars), significant main effects were found for pulse rate (*F*(1,7)=51.17, *p_corrected*=0.00019), and envelope type (*F*(5,35)=4.82, *p_corrected*=0.0167), and there was a significant interaction between these factors (F(5,35)=5.68, *p*=0.0056). In contrast, no significant main effects or interactions were observed for the ENV ITD probit coefficients (Fig. 4, red bars). In other words, PT ITD sensitivity depended on pulse rate and envelope type, but ENV ITD did not.

Post-hoc paired t-test comparisons to assess the effect of envelope width on PT ITD coefficients at 900 pps found significant differences between the 10 ms and the 50 ms wide envelopes (*p*=0.0106), as well as between the 10 and 200 ms wide envelopes (*p*=0.0315), suggesting that very rapidly rising envelopes facilitated the detection of PT ITDs. No significant pairwise differences as a function of envelope width were seen at 4500 pps (*p*>0.05), and no significant effect of whether the envelopes contained pauses was found irrespective of pulse rate (*p*>0.05).

One interesting detail one can observe in every panel of Figures 2 and 3 is that it is much easier to see an effect of ENV ITD when PT ITD is zero (in which case % “right” responses tend to increase from around 40% at −80 μs to around 60% at +80 μs) than when PT is not zero (in which case % right changes hardly at all as a function of ENV ITD. To test whether this tendency is statistically significant, we fitted that data shown in Figure 3 (pooled over all animals but separately for each pulse rate and envelope type) both with a simple Probit model as described above and an interaction model which allowed for ENV ITD coefficients to depend on whether PT ITD was zero or not (see Methods for details), and we used deviance tests to look for statistically significant differences in how well the models fit the data. In all cases the interaction model fitted significantly better (p values ranging from 0.008 to 2✕10^-9^) than the simple additive model. We also investigated whether the reduced effectiveness of ENV ITDs might be explained by “ceiling effects”. We know from previous studies (Li et al. 2019; Rosskothen-Kuhl et al. 2021; Buck et al. 2023) that the psychometric functions for ITD lateralization of rats are sigmoidal but may asymptote at “lapse rates” somewhere above 0% and below 100%, reflecting the fact that the animals will on some proportion of trials deviate from the optimal choice even if the stimuli are easy for them to discriminate. If we were to assume that +/-80 μs PT ITDs on their own provided a strong enough cue to push an animal’s choices very close to their performance ceiling (that is, the animal’s lapse rate for −80 μs PT ITD or 1-lapse rate for +80 μs PT ITD), then any additive effects of ENV ITD would become very difficult to discern since the animal is then operating in the “flat”, asymptotic part of its psychometric curve. To test whether ceiling effects are a plausible, sufficient explanation for the observation that ENV ITDs have a much smaller effect when PT ITDs are not zero, we fitted the data shown in Figures 2 and 3 with modified Probit models that incorporated a lapse rate as a free parameter (see Methods for details). Probit-with-lapse models gave a better fit to the data than simple Probit models, but they yielded often implausibly high lapse rate estimates, in about half the number of cases as high as 40%, well beyond the approximately 0-15% lapse rates we know to be typical from previous studies (Li et al. 2019; Rosskothen-Kuhl et al. 2021; Buck et al. 2023). Indeed, it is easy to see that the data shown in the left panel of Figure 2 is incompatible with the assumption that typical lapse rates for the animals in this study can exceed 15%. Given that our Probit-with-lapse model needed implausibly high lapse rates to produce a very good fit to the data suggests that ceiling effects are at best a partial explanation for the stronger effect of ENV ITD when PT ITD = 0 compared to when PT ITD ∈ {-80, +80} μs. We conclude that the observation that ENV ITD has a stronger effect if PT ITD is zero is statistically robust, but the cause of this non-additive interaction of ENV and PT ITDs remains unknown.

## Discussion

### PT ITDs are very powerful binaural cues in prosthetic hearing, but ENV ITDs are not

The key finding of this study is that PT ITDs are very powerful cues to sound location perception in the CI supplied mammalian pathway, while ENV ITDs are not. The difference in the effectiveness of these two cue types is very substantial: The mean PT ITD coefficient over all animals and stimulus conditions was 10.19, compared to a mean ENV ITD coefficient of 1.77 (compare height of blue and red bars in Fig. 4). To put these values in perspective, we can run our probit model (eqn. 1 below, see Methods) to predict how well an average rat with zero bias (*b_0_*=0) would lateralize either ENV or PT ITDs of a given magnitude. The model predicts that, when given informative PT ITDs of 0.1 ms at zero ENV ITD, it would lateralize these with an accuracy of 84.6% (a prediction that is in line with the values we observe in Figures 2 and 3), and informative PT ITDs of 0.3 ms would allow the animal to lateralize stimuli with 99.9% correct. In contrast, if we instead hold PT ITDs at zero but provide informative ENV ITD cues, the predicted performance would only reach an extremely modest 57% correct at 0.1 ms ENV ITD, and 70.2% at 0.3 ms. So even quite large ENV ITDs of 0.3 ms, which NH humans can lateralize with great ease, are barely sufficient to reach the ∼75% performance commonly considered a “just noticeable difference” threshold when delivered via CIs. These example calculations clearly illustrate that PT ITDs powerfully inform the ITD perception of our neonatally deafened biCI rats, while ENV ITDs make an in comparison almost negligibly small contribution. If PT ITDs and ENV ITDs conflict, as they routinely will in human patients fitted with contemporary clinical processors, the uninformative PT ITDs would be expected to dominate completely, which will disrupt the patient’s ITD perception. It is reasonable to assume that the patients’ brains will adapt to this confusing input by becoming desensitized to ITD, and this provides a highly likely explanation for the poor ITD sensitivity currently observed in human CI patients (Litovsky 2010; Ehlers et al. 2017).

### ENV ITD coding in normal hearing is unrepresentative of the CI case

As we have just seen, in the CI stimulated pathway, ENV and PT ITDs can easily confound each other, with PT ITDs playing a dominant role. It is important to appreciate that, in natural hearing, this type of confounding is much less of a problem because envelope and fine structure information become segregated already at the transduction stage in a manner that has no equivalent in bionic hearing with pulsatile stimuli.

We observed only a very modest sensitivity to ENV ITDs in the CI stimulated animals, even when the stimuli provided had very fast pulse rates (4500 pps) and steeply rising envelopes. Even though we stimulated the middle turn of the cochlea, which in normal, acoustic hearing, would provide inputs to high frequency “envelope”, rather than low-frequency “fine-structure” (FS), ITD processing pathways. The role of ENV ITDs in the CI-stimulated auditory system is therefore clearly very different from that observed in healthy, acoustically-stimulated ears. The likely reasons for these differences may become clearer if we quickly review how the NH cochlea encodes FS and ENV temporal information, respectively. Bear in mind that the FS versus ENV ITD distinction has nothing to do with ITD as an acoustic cue: in the free field, sound takes longer to travel to the farther ear, and this will delay all sound features, including fine structure and sound envelope, at the far ear by the same amount. The distinction of FS versus ENV ITD classically distinguishes how low or high frequency temporal features of a sound wave are encoded in the timing of AN action potentials. For frequencies below ∼1.5 kHz, the membrane potentials of the cochlear hair cells become entrained to the acoustic stimulus waveform. Consequently, the activity of AN synapses, and therefore the likelihood of AN action potentials, also become “phase locked” to peaks in the acoustic waveform. The differences in action potential timing in the left and right ears can then be compared to extract FS ITDs. However, if the frequencies of the sound waves increase beyond ∼1.5 kHz, hair cell membrane voltages can no longer follow each cycle of the acoustic waveform, and they then transition from an “AC” mode to a “DC” mode, in which their membrane voltage follows rises and falls in lockstep with the amplitude envelope of the sound, rather than the sound fine structure (Palmer and Russell 1986). At high frequencies, the firing of AN fibers therefore phase locks to envelope features (Frisina 2001). NH listeners will therefore detect FS ITDs in low sound frequencies (Zwislocki and Feldman 1956; Wightman and Kistler 1992; Macpherson and Middlebrooks 2002; Brughera et al. 2013), and ENV ITDs in high sound frequencies (Henning 1974; McFadden and Pasanen 1976) for the simple reason that sound transduction by cochlear hair cells performs an envelope extraction step if the sound frequencies exceed a relatively low phase locking limit.

Crucially, patients who require CI surgery almost invariably lack functioning cochlear hair cells. In any event, CIs completely bypass the envelope extraction mechanism that would normally occur in cochlear hair cells, and they directly trigger AN fiber discharges with extremely brief electric pulses. These electric pulses occur at fixed pulse rates, and spike times in the AN “snap to a temporal grid” laid down by the fixed rate pulse train because the probability of action potential initiation immediately after a pulse is very much higher than the probability of an action potential being initiated half way between two pulses. But if spike timing in each ear is constrained by the temporal grids laid down by the CI processors driving each ear, then we cannot expect inter-aural spike time differences to be able to reliably and accurately represent ITDs at a resolution that is much smaller than the “spacing” of the temporal grids, that is, the inter-pulse interval. Small ENV ITDs can therefore only be accurately represented in the auditory pathway of biCI patients if an as yet unknown physiological mechanisms can interpolate envelope features between “grid points”, and this envelope reconstruction would have to occur after the electrode-AN interface, but before the ITD processing stages of the superior olive. Data published in Figure 8 of Hancock et al. (2017) suggests that the interpolated envelope extraction that would be required in the early stages of the CI stimulated auditory brainstem to make resolving very fine ENV ITDs possible certainly does not occur routinely, if it occurs at all. In that study, the authors performed recordings in the IC of deafened, cochlear implanted cats in response to amplitude modulated pulse trains. They were able to show that the spike timing of these IC neurons, which the authors describe as typical, locked to the first few supra-threshold pulses that sampled the envelope. Thus, at the level of the auditory midbrain, well past the point where ITDs are normally extracted, the timing of spikes still typically “snaps to the temporal grid” laid down by the electrical stimulus pulse train. This implies that no “reconstruction filter” has been applied that would interpolate spike times between grid points and thereby resolve temporal envelope features at a resolution finer than the inter-pulse interval (about 1 ms at the ∼1000 pps pulse rate typically used in clinical processors).

A complete failure of envelope reconstruction prior to ITD computation would lead to an absolute dominance of PT ITDs over ENV ITDs in ITD lateralization judgments. Conversely, perfect envelope reconstruction would lead to ENV ITDs completely dominating over PT ITDs. Our data clearly indicate a strong dominance of PT ITD over ENV ITD, but ENV ITDs nevertheless had a small influence. The influence of the temporal envelope on ITD judgments is almost negligibly weak compared to that of pulse timing, which suggests that the neural representation of temporal envelopes in the early stages of the binaural integration pathways of the CI stimulated auditory brainstem must be in some way weak or incomplete, even if, intriguingly, it appears not to be entirely absent. However, if it was possible for temporally very precise envelope representations to be strengthened by experience and to become independent of the neural representation of pulse timing, then we would expect human biCI users to be able to develop ENV ITD thresholds that should be comparable to those seen in NH listeners. A wealth of data on biCI patients indicates that this is not the case (van Hoesel and Tyler 2003; Laback et al. 2004; Majdak et al. 2006; Noel and Eddington 2013).

Note also that the loss of envelope extraction that would normally occur at the cochlear hair cells for high-frequency inputs also implies that the large body of psychoacoustic studies of ENV ITD sensitivity in NH individuals is not directly applicable to the biCI case, because no equivalence can be assumed between the neural representations of acoustic sound stimulus envelopes and electric pulse train stimulus envelopes in the early stages of the auditory pathway.

### Comparison to previous work on human biCI patients

As mentioned in the introduction, a key motivation for the current study was to use an animal model to overcome confounds, which arise from the fact that human patient data is invariably collected against a background of typically many months of daily prior stimulation from clinical processors delivering misleading PT ITD cues. With this point in mind, it is instructive to compare our results against previous studies on human patients, and of particular interest is a paper by Noel and Eddington (2013), which used experimental processors to measure PT ITD and ENV ITD thresholds in six biCI patients. Although there are some differences between their approach to ours (they opted to run adaptive staircases in which they varied either PT ITD or ENV ITD while holding the other cue constant at zero, or they co-varied both in lock-step), their approach was in many ways so similar to ours to make a direct comparison very illuminating: like us, as they tested sinusoidal envelopes of varying widths imposed on 1000 pps pulse train carriers, quite similar to the 900 pps trains we used here. But while their methods were comparable, their findings are in very important respects radically different.

Firstly, 5 out of 6 of their human patients had PT (“carrier only”) ITD thresholds too large to measure (in this case greater than 450 μs). Only one participant had a measurable 71% PT ITD threshold of 133 μs. Contrast this with our rat results, where all 8 of our animals performed at over 80% correct for PT ITDs of 80 μs (see Fig. 3, top row). In fact, we deliberately tested only +/-80 μs ITD values because our previous work has shown that this is supra-threshold for biCI stimulated rats (Rosskothen-Kuhl et al. 2021; Buck et al. 2023). The PT ITD threshold of our rats is reliably and dramatically better than that seen in the human patients tested by Noel and Eddington (2013). Curiously, even though 5 out of 6 of their patients showed no measurable sensitivity to PT ITD alone, PT ITDs nevertheless showed a synergistic effect in the sense that stimuli in which ENV and PT ITDs agreed produced markedly lower thresholds than stimuli where PT ITDs were held at zero and only ENV ITDs were varied. This result is broadly in line with observations by Churchill et al. (2014) who observed beneficial effects of providing PT ITDs in a sound localization task on biCI patients. Nevertheless, the effect of PT ITD in the Noel and Eddington (2013) study was comparatively small, as is so often the case in studies on human patients, while in our rats the effect of PT ITDs was clearly and powerfully dominant.

Secondly, ENV ITD thresholds of most patients were in most cases substantially better than the very poor PT ITD performance, with typical 71% correct thresholds between 120 and 300 μs (Noel and Eddington 2013). Laback et al. (2004) reported similar ENV ITD thresholds of 259 and 384 μs, respectively, from two patients tested with clinical processors. While our study was not designed to measure ENV ITD thresholds directly, as mentioned above, we can extrapolate the probit model fit to our data to estimate 70% correct thresholds to be about 300 μs for our biCI rats. Extrapolations always need to be considered low confidence estimates, but it is interesting that the extrapolated ENV ITD thresholds for our rats turn out very similar to the values observed by Noel and Eddington in their human patients. van Hoesel and Tyler (2003) report an ENV ITD threshold of 290 μs for a single biCI patient, which is also in a similar range.

Noel and Eddington (2013) results do contrast somewhat with an earlier study by Majdak et al. (2006), which also investigated ITD sensitivity in biCI patients with amplitude modulated stimuli, and which did report PT ITD sensitivity in four out of four biCI patients tested, but typically only at much lower pulse rates than the 900 to 1000 pps typically used either in clinical settings, or in this study and that by Noel and Eddington (2013). Majdak et al. (2006) also only tested ITDs in very large steps, and their methodology does not lend itself to direct comparisons with the responses to the much smaller ITDs used in our study. We note, however, that Majdak et al. (2006) observed a significant, direct effect of ENV ITD in only one out of four CI patients, so in their study as in ours, ENV ITD turned out to be a much weaker cue than PT ITD. Noel and Eddington (2013) patients had used bilateral clinical processors for between 10 months and 6 years. Interestingly, the one single patient who exhibited a measurable PT ITD threshold at 1000 pps had the second lowest amount of experience listening with bilateral clinical CIs (1 year), much less than the median experience of 3 years.

Our results and all previous studies we are aware of are compatible with the idea that chronic stimulation with clinical biCIs, which present confusing PT ITDs over a period of many months, may lead to a loss of sensitivity to PT ITDs that would otherwise be a very powerful lateralization cue. Also supportive of this notion are recent reports that pre-lingually deafened patients fitted with clinical processors, such as MED-EL devices implementing the FS4 strategies, which do encode at least some temporal FS information in the pulse timing delivered to low frequency channels, are more likely to exhibit ITD thresholds well below 1000 μs when tested with low frequency sounds than patients who received no pulse-timing encoded FS information at all (Eklöf and Tideholm 2018). Similarly, Fischer et al. (2021) report improved ITD thresholds when post-lingually deafened biCI patients are stimulated with FS4 strategies as opposed to alternative strategies which do not encode ITDs in pulse timing.

However, another recent paper by Ausili et al. (2020), which compared patients with or without FS encoding, found that most of their patients had very poor ITD sensitivity irrespective of their FS status with no statistically significant difference between groups, but note that the patients tested in that study were a very heterogenous group, with ages ranging from 22 to 77 years, and greatly varying ranges of experience with the biCI devices. Also, almost all the patients in the Ausili et al. (2020) study underwent periods of between 1 and 7 years during which they were only monaurally stimulated - a situation which is bound to be unfavorable to the development of precise bionic binaural hearing (Gordon et al. 2014; Killan et al. 2019). The great heterogeneity of the cohort tested by Ausili et al. (2020) most likely explains the very large intra-subject variability in their dataset. Eklöf and Tideholm (2018), in contrast, studied a more homogenous cohort of young, early implanted participants. Such discrepancies in the literature, and the often unavoidable confounds that can arise when working with heterogenous patient populations, underscore the value of experimental animal studies in which variables of interest can be much more precisely controlled and isolated from confounding variables.

### Effects of pulse rate and envelope width and rate

For the relatively small ITDs we focussed on in this study, we observed significant effects of envelope width (or, equivalently, modulation rate) and pulse rate only on PT ITD sensitivity, but not on ENV ITD sensitivity. The effects on PT ITD sensitivity (greater sensitivity at 900 than 4500 pps and for more rapidly, rather than slowly, rising envelopes) are entirely in line with expectations from previously published studies (van Hoesel and Tyler 2003; Litovsky 2005; van Hoesel et al. 2009; Buck et al. 2023). ENV ITD sensitivity was uniformly low and similar for all envelope widths, envelope rates, and pulse rates tested.

In our experiments, we also observed no significant change in lateralization performance as a function of whether the envelopes included silent pauses or not. While some researchers have suggested that silent pauses may have a positive effect on ITD sensitivity in normal hearing (Klein-Hennig et al. 2011; Greenberg et al. 2017) and prosthetic hearing (Hancock et al. 2017), our results do not confirm this. Instead, they are in agreement with another set of studies by Laback and Majdak (2008); Noel and Eddington (2013) and Goupell et al. (2009) who also did not find an effect of pauses in their assessments of binaural hearing in CI patients.

Additionally, our findings that a sharper onset facilitates the detection of ITDs are in agreement with Greenberg et al. (2017) who found better ITD sensitivity for smaller burst width in IC recordings of guinea pigs. Similar observations were made by Noel and Eddington (2013) and Klein-Hennig et al. (2011) who found better ITD sensitivity for sharper onsets in their CI patient and NH subjects, respectively. However, we only observed significant effects for PT ITD at 900 pps.

### Comparison to previous electrophysiological work in CI animals

Smith and Delgutte (2008) previously attempted to reveal the possible effect of ENV ITDs on the biCI stimulated nervous system using an electrophysiological approach. They anesthetized hearing experienced cats, implanted their inner ears bilaterally, and then recorded the responses of inferior colliculus (IC) neurons to binaural 1000 pps or 5000 pps pulse trains with fixed PT ITD but continuously, slowly varying ENV ITD. The rationale behind this approach was that continuously varying ITDs may create “binaural beats” (https://auditoryneuroscience.com/spatial-hearing/binaural-beats) that can generate a vague sense of movement in a listener, and that should produce modulations in the firing rates of ITD sensitive neurons in the auditory pathway that are time locked to the rate of ITD variation. Smith and Delgutte (2008) did indeed observe such firing rate modulations that occurred in step with ENV ITD modulations, which led them to conclude that there must be at least some sensitivity to ENV ITDs in the auditory pathway of biCI stimulated cats. Their experimental approach was quite imaginative, but it is perhaps hard to map onto conventional listening conditions (binaural beat-like, gradual ITD variations in an otherwise constant sound are very rare in natural or urban auditory environments). Additionally, the very large range and large step size of ENV ITDs used in their study does not lend itself to an accurate estimation of neural ENV ITD thresholds. Nevertheless, by comparing the effect of ENV ITD modulation to that of changing PT ITDs in fixed step sizes, Smith and Delgutte (2008) concluded that the biCI stimulated mammalian auditory pathway appears to be much more sensitive to moderately small steps in PT ITD than in ENV ITD, and that these effects are observable at the relatively high pulse rates typically used in clinical settings (not just the 300 pps or less so often used in behavioral tests on human patients). Extrapolating from observed firing rate changes recorded in the auditory midbrain of an anesthetized animal to actual perceptual capacities of an awake, listening participant is of course a difficult exercise (Parker and Newsome 1998). Our behavior experiments described here, conducted in a very different mammalian species and using a very different approach, nevertheless agree very well with what one might have expected to see based on their data, and they strongly confirm the validity of their conclusion “that a bilateral cochlear implant strategy that successfully conveys ITD cues should control the precise timing of current pulses based on the fine timing of the sound sources at each ear”. Our data provide the most direct and compelling evidence yet that ITDs encoded in pulse train envelopes are simply not up to the task of eliciting a sensory sensitivity to the very subtle cues required to provide CI patients with binaural listening experiences that are as rich, detailed, and highly informative as those that we normal hearing listeners enjoy and too easily take for granted.

## 4. Conclusion

In normal hearing, even very small ITDs in the order of 100 μs or less provide powerful cues to sound source location. Our data show that, via biCIs, such small ITDs can only be delivered through pulse timing, not in signal envelopes, and this is true irrespective of envelope shape. Furthermore, in the recently implanted auditory system, PT ITDs are so much more powerful than ENV ITDs that the inappropriate PT ITDs typically delivered by devices in current clinical use will swamp the effect of other, potentially useful binaural cues. Most processors in clinical use make no effort at all to represent temporal features of the auditory input in the timing of stimulus pulses with the required precision of a small fraction of a millisecond, and the only devices that do offer a degree of pulse time encoding, such as the FS4 strategy by MED-EL, do so only for a small subset of the cochlear stimulation channels. In the light of our results presented here, we hypothesize that the almost complete absence of accurately timed stimulus pulses in the input delivered to biCI patients is the primary driver of poor binaural hearing outcomes in this patient population.

## Materials and Methods

All procedures involving experimental animals reported here were performed under license issued by the Department of Health of Hong Kong (#16-52 DH/HA&P/8/2/5) and the Regierungspräsidium Freiburg (#35-9185.81/G-17/124). They were also approved by the City University of Hong Kong Animal Research Ethics Sub-Committee. A total of 8 female Wistar rats were used in this study. All rats were neonatally deafened and underwent acoustic and electric auditory brainstem response (ABR/eABR) recordings, bilateral cochlear implantation and behavioral training as described previously (Rosskothen-Kuhl et al. 2021; Buck et al. 2023) and here below.

### Neonatal deafening and cochlear implantation

All rats were neonatally deafened by daily injections of kanamycin from postnatal day 9 to 20 inclusively as described in Rosskothen-Kuhl et al. (2021). This is known to cause widespread death of inner and outer hair cells while keeping the number of spiral ganglion cells comparable to that in untreated control rats (Osako et al. 1979; Matsuda et al. 1999; Argence et al. 2008). Profound hearing loss (>90 dB) was confirmed by the loss of Preyer’s reflex (Jero et al. 2001) and the absence of auditory brainstem responses (ABRs) to broadband click stimuli and pure tones (500, 1000, 2000 and 8000 Hz). For a subset of the animals, the absence of hair cells in the cochlea was also histologically confirmed.

The animals were raised to young adulthood (postnatal day 70 to 119) and then implanted simultaneously with biCIs under ketamine (80 mg/kg) and xylazine (12 mg/kg) anesthesia. The electrodes were inserted via a cochleostomy over the middle turn, and their function was confirmed using eABRs.

### Electric stimulation

The electric stimuli used to examine the animals’ eABR and behavioral ITD sensitivity were generated using a Tucker-Davis Technology (TDT, Alachua, FL) IZ2MH programmable constant current stimulator at a sample rate of 48,828.125 Hz. One of the tip CI electrodes served as the stimulating electrode, the adjacent electrode as ground electrode. All electrical intracochlear stimulation used biphasic current pulses similar to those used in clinical devices (duty cycle: 40.96 ms positive, 40.96 ms at zero, 40.96 ms negative), with peak amplitudes of up to 300 mA, depending on eABR or informally assessed behavioral comfort levels (rats will scratch their ears frequently, startle or show other signs of discomfort if stimuli are too intense). For behavioral training all neonatally deafened biCi rats were stimulated on average at levels ranging from 3-8 dB above these eABR thresholds, depending on the pulse rate, modulation rate, and repetition rate. Careful observation of the animals’ behavior during spontaneous presentations of test stimuli was used to confirm that stimulus levels were high enough to be reliably detected, but not so high as to cause discomfort. Behavioral stimuli from the TDT IZ2MH were delivered directly to the animal through a custom built head connector that was connected and disconnected before and after each training session. All biCI rats received binaurally synchronized input from the first stimulation. For full details on the electric stimuli and stimulation setup see Rosskothen-Kuhl et al. (2021).

### Psychoacoustic training and testing

Following implantation, animals were trained on a two-alternative forced-choice ITD lateralization task in our custom made behavior setup as described in Rosskothen-Kuhl et al. (2021); Buck et al. (2023). The behavioral procedure is illustrated in Figure 1A. Briefly, this setup contained three water spouts from which the animal could receive water as reward. The animal was trained to lick the center spout to start a trial and trigger the delivery of a biCI stimulus pulse train. The animal would then need to make a behavioral choice and lick either the left or right spout to indicate on which side they heard the stimulus. Correct responses were rewarded with drinking water, incorrect responses received negative feedback in the form of a short timeout.

For the initial training PT ITDs and ENV ITDs always co-varied, as they would in natural hearing, and they varied over a range of +/-150 µs, where negative ITD values denote left leading, positive values right leading ITDs. Previous studies had shown that rats can lateralize +/-80 µs ITDs reliably and easily. Training concluded, and formal testing commenced, once animals were able to lateralize ITDs of 80 µs or greater highly reliably (>75% correct). The animals would typically reach this criterion performance within two weeks of training. Some of the animals in this study took part in another ITD lateralization task with strictly co-varying ENV and PT ITDs prior to taking part in this experiment (Buck et al. 2023), and they required essentially no additional training. In the subsequent testing sessions, ENV ITDs and PT ITDs were allowed to vary independently from the set {-80, 0, +80} µs, under the constraints that ENV ITDs and PT ITDs could not both be zero. Thus, there were effectively four different ITD configurations: “congruent”, with ENV ITD and PT ITD pointing in the same direction, “incongruent”, with ENV ITD and PT ITD pointing in opposite directions, “PT only”, in which only the PT ITD differed from zero, and “ENV only”, in which only the ENV ITD differed from zero. In order not to influence the animals’ use of ENV or PT ITDs in their lateralization judgments through the reward schedule, we rewarded the animals whenever the side on which they responded corresponded to the side on which either the PT ITD or the ENV ITD were leading. This meant that the animal had a free choice on incongruent trials and would be rewarded regardless. The four different ITD types were presented in random order, such that ⅓ of trials were congruent, ⅓ PT only, ⅙ ENV only and ⅙ incongruent. The frequency of incongruent trials was reduced to keep the proportion of trials in which the animals might be able to get a reward even if they weren’t paying attention small. The frequency of ENV only trials was reduced because informal piloting had suggested that lateralizing ENV only trials is very hard for the animals, and to keep the animals motivated to continue paying attention to the stimuli, it was necessary to keep the proportion of trials in which this attention leads to reward relatively high. The carrier pulse rates for the testing stimuli were chosen pseudorandomly for each trial to be either 900 or 4500 pps. Different stimulus envelope types were tested, differing in envelope widths (10, 50, 200 ms) and rates (1, 5, 20, 100 bps) with a modulation depth of 100% as illustrated in Figure 1 C. But to make it easy for the animals to focus on lateralizing ITDs without the distraction that would arise from constant and very wide variation in envelope types, only one or at most two envelope types were tested during any one of an animal’s two daily testing sessions.

In a typical session, a rat performed approximately 200 trials, or about 400 trials daily, five days a week, with rest and ad lib water on weekends. In total, the rats in this study performed between 6500 and 12600 trials in the course of between 42 and 65 sessions. In total, the data set for this study comprises 73484 trials.

### Probit regression

To quantify the effects of PT ITD and ENV ITD illustrated in Figure 2, we fitted the behavioral data with probit regression models of the form:

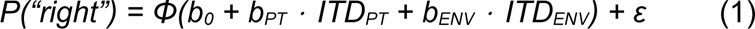

where *Φ* is the cumulative gaussian function, *ITD_PT_* and *ITD_ENV_* are the respective ITD values in ms {-0.08, 0, +0.08}, *b_PT_* and *b_ENV_* are the coefficients (“regression slopes”) that capture the animal’s sensitivity to PT ITD and ENV ITDs, respectively, *b_0_* is a constant parameter which can account for any small, idiosyncratic preferences (biases) an animal may have for one side over the other, and ε is the residual error. The model parameters were fitted using the function *discrete.discrete_model.Probit()* from the *statsmodels* Python library (https://www.statsmodels.org/) as detailed in the shared code accompanying this manuscript (see below).

This probit model assumes that the z-value of a “right” response increases linearly with ITD, a reasonable assumption given that psychometric functions derived from cumulative gaussians often provide a good fit to rat ITD lateralization data (Li et al. 2019). The coefficients *b_PT_* and *b_ENV_* thus quantify the behavioral sensitivity to PT ITDs and ENV ITDs in a manner that is directly comparable and fairly easily interpretable (see Fig. 4).

To test for non-additive interactions between PT ITD and ENV ITD and to investigate whether these interactions might be explained by a ceiling effect we adapted our analysis code from Li et al. (2019), as this allows us greater freedom in fitting richer model equations. That code uses gradient-descent to discover maximum likelihood parameters for fitting model equations to our data by. To ask whether the apparent differences in ENV ITD effects as a function of PT ITD were statistically significant, we compared the fits of two models, one which allows only a single *b_ENV_*parameter to capture the effect of ENV ITD, and a second which allows two.

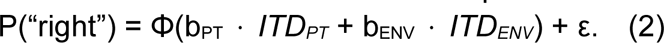

which is the same as in eqn 1, but it omits the bias term b0, which can be easily done as biases in the data were small throughout. This was compared against an interaction model

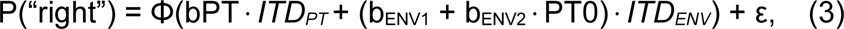

which introduces two separate ENV ITD weights (b_ENV1_ and b_ENV2_) as well as an indicator variable PT0, which takes the value of 1 if ITD_PT_=0, or 0 otherwise. This model will thus fit two different Probit regression slopes for ENV ITD: b_ENV1_ for non-zero PT ITD conditions, and (b_ENV1_ + b_ENV2_) for zero PT ITD conditions. To test which of these models fit the data better we conducted deviance tests, which take into account that the difference in deviances between two generalized linear models is χ^2^ distributed with a number of degrees of freedom equal to the difference in the number of free parameters of the models. As described in the Results, the interaction model fitted our data highly significantly better.

To assess the possibility that ceiling effects might explain the significant non-additive interactions between ENV ITD and PT ITD, we attempted to fit our data with a model similar to that developed in Li et al. (2019), which included a lapse rate parameter. The model equation was

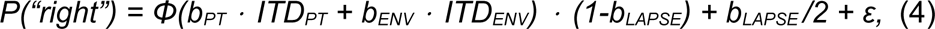

where *b_LAPSE_* was the lapse rate parameter, a real number in the interval [0,0.5] that captures the proportion of trials in which the animal is thought to make errors that are unrelated to its underlying ability to discriminate the stimuli correctly. As mentioned in the Results section, the maximum likelihood fits of this model to our data often required implausibly high estimates for the value of *b_LAPSE_*in order to produce a very good fit to the data, indicating that ceiling effects are likely insufficient to explain the observed interactions between ENV ITD and PT ITD.

## Data Access

All data as well as the analysis code used to generate all the figures and statistical results included in this manuscript can be downloaded from https://auditoryneuroscience.org/dataShare.

## Acknowledgments

The work leading to this publication was supported by the German Academic Exchange Service (DAAD) with funds from the German Federal Ministry of Education and Research (BMBF) and the People Programme (Marie Curie Actions) of the European Union’s Seventh Framework Programme (FP7/2007-2013) under REA grant agreement n° 605728 (P.R.I.M.E. – Postdoctoral Researchers International Mobility Experience), as well as grants from the Hong Kong General Research Fund (11100219), the Hong Kong Health and Medical Research Fund (06172296), the Shenzhen Science Technology and Innovation Committee (JCYJ20180307124024360), the Martin Lee Centre for Innovations in Hearing Health at Macquarie University, the Research Commission of the Medical Faculty of the Medical Center at the University of Freiburg, and the charity “Taube Kinder lernen hören e. V.“. Eight cochlear implant animal arrays were kindly provided by MED-EL Medical Electronics, Innsbruck, Austria (Research Agreement PVFR2019/2).

The authors would also like to thank Stella Mayer for their assistance in collecting data.

